# Structural basis for NONO specific modification by the α-chloroacetamide compound *(R)*-SKBG-1

**DOI:** 10.1101/2025.06.10.658868

**Authors:** Alessia Vincenza Florio, Corinne Buré, Sébastien Fribourg

## Abstract

Among the many proteins involved in cancer progression an increasing number of RNA Binding Proteins (RBPs) are central to the function of a cell and tightly associated to genetic diseases as well as cancer appearance and progression. In a recent study, small molecule inhibitors have been identified as targeting NONO, a RBP known to be involved in mRNA splicing, DNA repair and membraneless organelles stability. Here we report the molecular basis of NONO-targeting by the α-chloroacetamide *(R)-*SKBG-1. We explore the specific binding and enantiomer specificity of NONO towards *(R)*-SKBG-1 using mass spectrometry and structure determination. We have determined the crystal structure of *(R*)-SKBG-1-bound to NONO homodimer. This study sheds light on the conformational plasticity of *(R)*-SKBG-1 when covalently bound to NONO. Altogether these results give an experimental rationale for ligand modification and optimization in a future use as a drug against cancer.

**SIGNIFICANCE:** DBHS proteins form a family of three proteins encoded by three different and essential genes. They form obligate homodimers and heterodimers to fulfil their function. In the cell, they are involved in mRNA splicing, DNA repair and membraneless organelles formation. Recently, NONO has been identified as a target of small-molecule inhibitors in prostate cancer cells. Treatment with α-chloroacetamide modifies a specific cysteine residue only found in NONO and not in its paralogue proteins SFPQ and PSPC1. Here we provide the molecular basis of α-chloroacetamide covalent binding to NONO and we explore the enantiomer specificity of binding. We also demonstrate that α-chloroacetamide can target NONO in homodimers and heterodimers and that both binding sites are equivalently modified. Finally, we provide show that α-chloroacetamide binding to NONO is driven by the combination of covalent binding and conformational flexibility of the ligand. Altogether, we believe that this study provides useful information for ligand improvement aiming at targeting NONO in cancer cells.

**OUTLINE:**

- NONO residue C145 is targeted by *(R)-*SKBG-1
- The two binding sites are equally modified in NONO homodimers
- NONO is specifically targeted and not SFPQ and PSPC1
- *(R)-*SKBG-1 adopts multiple conformations in the absence of RNA

## INTRODUCTION

RNA-binding proteins (RBPs) are critical effectors of gene expression through the regulation of processing, cellular localization, post-transcriptional modifications and mechanisms of degradation. As such, their malfunction underlies many diseases including rare genetic diseases and cancer ^1^. In most cases, their mode of RNA binding correlates with a protein surface that lacks traditional ligand binding pockets in comparison to enzymes and hormone receptors and therefore in spite of their fundamental roles, RBPs have been poorly explored as targets of chemical probes for diagnostics and therapeutics. Recently, Kathman et al. identified α-chloroacetamides as specific binders of NONO in a screen aimed at discovering small molecules that reduce the expression of transcripts encoding Androgen Receptor (AR) mRNA and its splice variants in prostate cancer (PCa) cells (Kathman et al, 2023).

Indeed, the Androgen Receptor (AR) signalling is critical in the development, function and homeostasis of the prostate. Hence it is not surprising that 80-90% of the PCa cases are dependent on androgens. In order to cure PCa, the first line of therapy aims at antagonizing the effect of androgen and inhibits its transcriptional activity. Despite an initial favourable response, almost all patients invariably progress to a more aggressive, and resistance to these drugs often emerges. To counteract this resistance, strategies aiming at degrading the AR using Proteolysis Targeting Chimeras (PROTACs) have been developed (Salami et al, 2018), but such strategies do not target splice variants for example.

NONO is one of the paralogues of the Drosophila/Behavior human Splicing (DBHS) protein family that comprises also SFPQ (Splicing Factor Proline and Glutamine Rich) and PSPC1 (Protein Splicing Component 1)^4,5^. All are essential proteins and involved in either transcriptional and post-transcriptional events including pre-mRNA splicing, DNA repair, circadian rhythm ^4^. At the molecular level, the DBHS proteins form obligate dimers, either as homodimers or heterodimers. The structure of the six dimer combinations have been obtained revealing an arrangement in a head-to-tail fashion in which the core-folded domain makes extensive contacts to form the dimer interface ^6–11^. No structural data is available for the DBHS proteins N-terminal and C-terminal non conserved extensions that are believed to play a function *in vivo* in modulating protein aggregation into liquid-liquid phase separation ^12,13^.

In their initial report, Kathman et al. could not address a number of points regarding NONO targeting by *(R)*-SBKG-1 ^2^. Indeed, as NONO not only homodimerizes but also heterodimerizes with SFPQ and PSPC1, it was not known whether NONO could also be targeted by α-chloroacetamide in heterodimers. It was not known whether both sites in NONO homodimer could be modified at the same time and to the same extent. Finally, the rationale for the observed *(R)* over *(S)* enantiomer selectivity could not be addressed in the study. In order to tackle these questions, we performed biophysical and structural studies on NONO qnd DBHS proteins after modification with the active and the inactive enantiomers of SKBG-1. Here we report the molecular basis of NONO interaction with the α-chloroacetamide *(R)*-SKBG-1 validated by mass spectrometry. We present the crystal structure of *(R)-*SKBG-1-bound to NONO homodimers.

## RESULTS

### NONO is efficiently modified by *(R)*-SKBG-1 on C145

In order to detect chemical modification of NONO after treatment with *(R)* & *(S)* SKBG-1 enantiomers, we used mass spectrometry under denaturing conditions (ESI-MS). The first series of experiments aimed at determining the optimal sample treatment conditions in order to measure with accuracy the molecular mass of NONO’s construct. To better visualize the effect of the treatment with (*R*) and (*S*) SKBG-1 enantiomers, we have shown a part of mass spectra in Figures corresponding to one charge state (in this case, z = 23), the entire mass spectra being shown in the Supplemental Figures. Experimental masses of the proteins were obtained by considering all the charge states of the ionized protein. As shown in Fig. 1A (and Fig. S1), we determined the mass after TEV cleavage of NONO(53-312) at M= 30,425.2 ± 0.7 Da (for m/z conversion to a mass please refer to Table S1) which corresponded to the expected mass (Table S1). We next explored the chemical modification of NONO after incubation with the active or the inactive chemical compounds *(R)*-SKBG-1 and *(S)-*SKBG-1 respectively, as defined by Kathman and colleagues ^2^. In NONO construct, two potential sites could be modified at position C145 and C208. Protein samples were incubated at 25 °C for 2h with a molecular ratio of ligand/protein of 2:1. A control sample followed the same incubation but without ligand. After treatment, the samples were extensively buffer exchanged by iterative concentration and dilution steps with 500 mM ammonium acetate at pH 7.0. As shown in Fig. 1B (and Fig. S1B)), a new peak appeared on the mass spectrum at a mass of M = 30,884.5 Da. This increase of mass of 459.3 Da (theoretical M=459.5) corresponded to a covalent adduct of *(R)-*SBKG-1 to NONO. Indeed, during the SN2 chemical reaction between NONO and the *(R)-* SBKG-1, the chlorin atom from the ligand is lost in addition to a H atom from the SH-of the C145 (Fig. S2A) resulting in a theorical addition of mass to NONO of 459.5 Da. Trace amounts of unmodified NONO can be observed and no secondary modification site is detected. Treatment with *(S)-*SKBG-1 shows poor modification of NONO under the same conditions (Fig. 1C). These data demonstrate a strong enantiomer preference for *(R)-*SKBG1-1 over the *(S)-* SKBG-1 under our experimental conditions and a full capacity of the *(R)-*SKBG-1 compound to react with every NONO molecule. This also reveals that only one site at a time is present in NONO construct used in this study.

**Figure 1:**
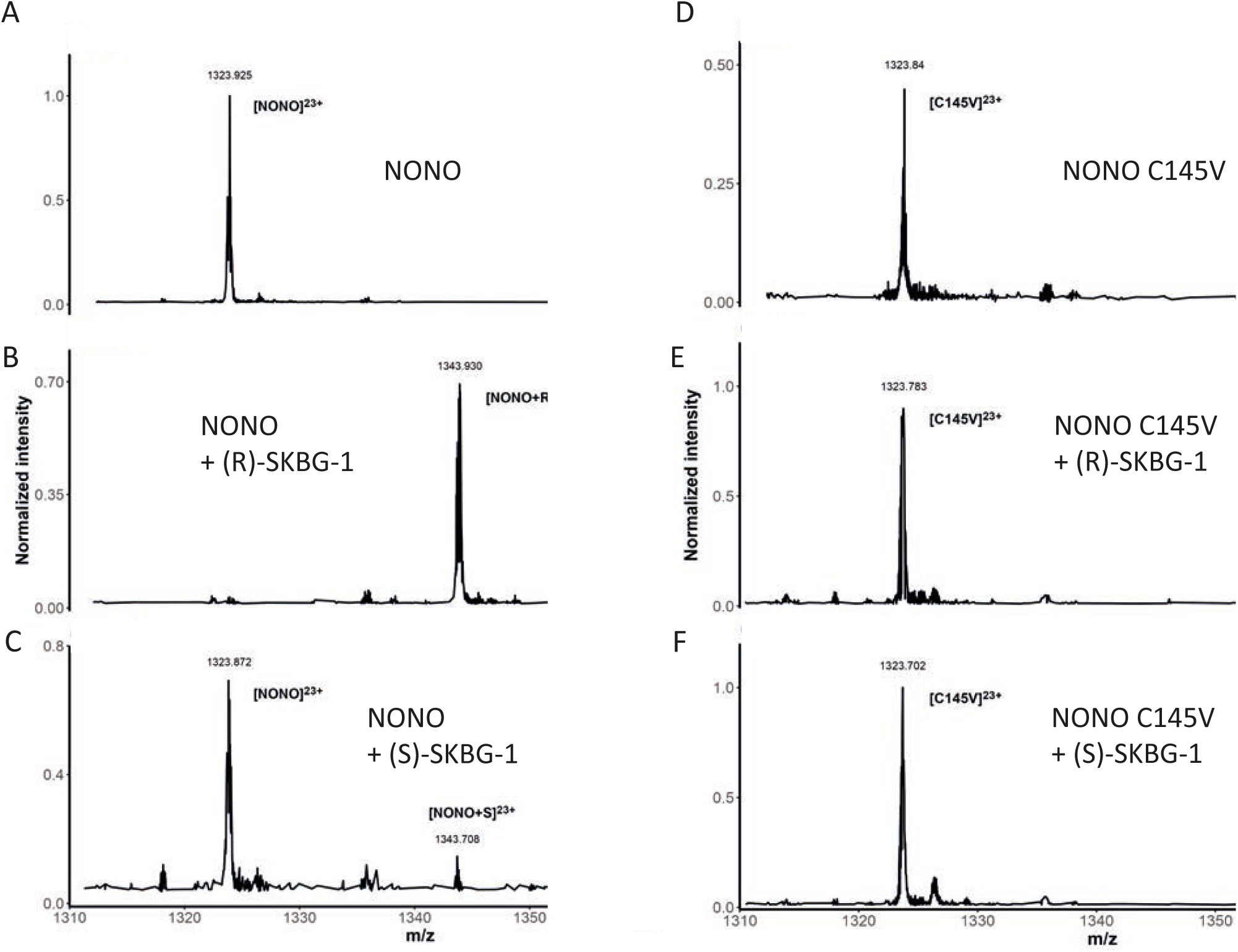
Mass determination by ESI-MS of native NONO and NONO C145V mutant before and after treatment with α-chloroacetamide. A) ESI-MS measurement of NONO(53-312). Here only the most intense charge state (z= +23) is shown for clarity. The ion [NONO+23H]^23+^ is annotated as [NONO]^23+^ also for clarity. This applies for for all panels. B) ESI-MS measurement of NONO(53-312) after treatment with *(R)-*SKBG-1. A peak corresponding to an increase of mass of 459.3 Da is measured. C) ESI-MS measurement of NONO(53-312) after treatment with *(S)-*SKBG-1. Only limited modification by (S)-SKG-1 is observed. D) NONO C145V point mutant sample was analyzed by ESI-MS. E) after treatment with *(R)*-SKBGR-1 and with *(S)-*SKBG-1 (F). No difference in spectra can be observed indicating that the point mutant C145V prevents ligand binding. Full spectra are shown in Fig. S1 and Fig. S2.

We next challenged NONO’s ligand-binding site. To this end we performed similar experiments as described above but with the C145V point mutant of NONO(53-312) (Fig. 1D and Fig. S3). No mass change could be detected after incubation with *(R)-*SKBGR-1 (Fig. 1E) nor *(S)-*SKBGR-1 (Fig. 1F) in comparison to the wild type protein (Fig. 1D). This demonstrates that C145 is indeed the only site of modification for the α-chloroacetamide as described by Kathman et al. ^2^, it also reveals that each binding site at C145 is equivalently accessible and modified in NONO homodimers.

### NONO is specifically modified in heterodimers

Having demonstrated that NONO(53-312) can be covalently modified by *(R)-*SKBG-1 in NONO homodimers, we next explored the modification of NONO in NONO-SFPQ and NONO-PSPC1 heterodimers. We applied the same experimental protocol as described above and we measured the mass of unmodified and modified complexes after incubation with *(R)-*SKBG-1 and *(S)-*SKBG-1 (data not shown). As shown on Fig. 2A (and Fig. S4, Table S1), the native form of NONO can be observed at a M= 30,231.2 Da (m/z of 1315.58 corresponded to [NONO+23H]^23+^), we could also accurately measure the mass of ligand-bound NONO(53-312) at a M= 30,691.4 Da (m/z of 1335.419 corresponded to [NONO+R+23H]^23+^). Moreover, only the mass of SFPQ(276-535) at M= 30,115,1 Da (m/z = 1310.361 for [SFPQ+23H]^23+^)(Fig. 2A and 2B) and PSPC1(62-320) M= 30,092,2 Da (m/z = 1309.273 for [PSPC1+23H]^23+^) (Fig. 2C and 2D) could be measured indicating that *(R)*-SKBG-1 specifically reacts with NONO and not with SFPQ nor PSPC1. We did not detect any modification of either NONO, nor PSPC1 nor SFPQ after treatment by *(S)*-SKBG-1 (not shown). Altogether, these experiments demonstrate that *(R)*-SKBG-1 reacts with NONO regardless of the complex it is involved in and despite the presence of multiple several cysteine residues in SFPQ (2) and PSPC1 (2) constructs.

**Figure 2:**
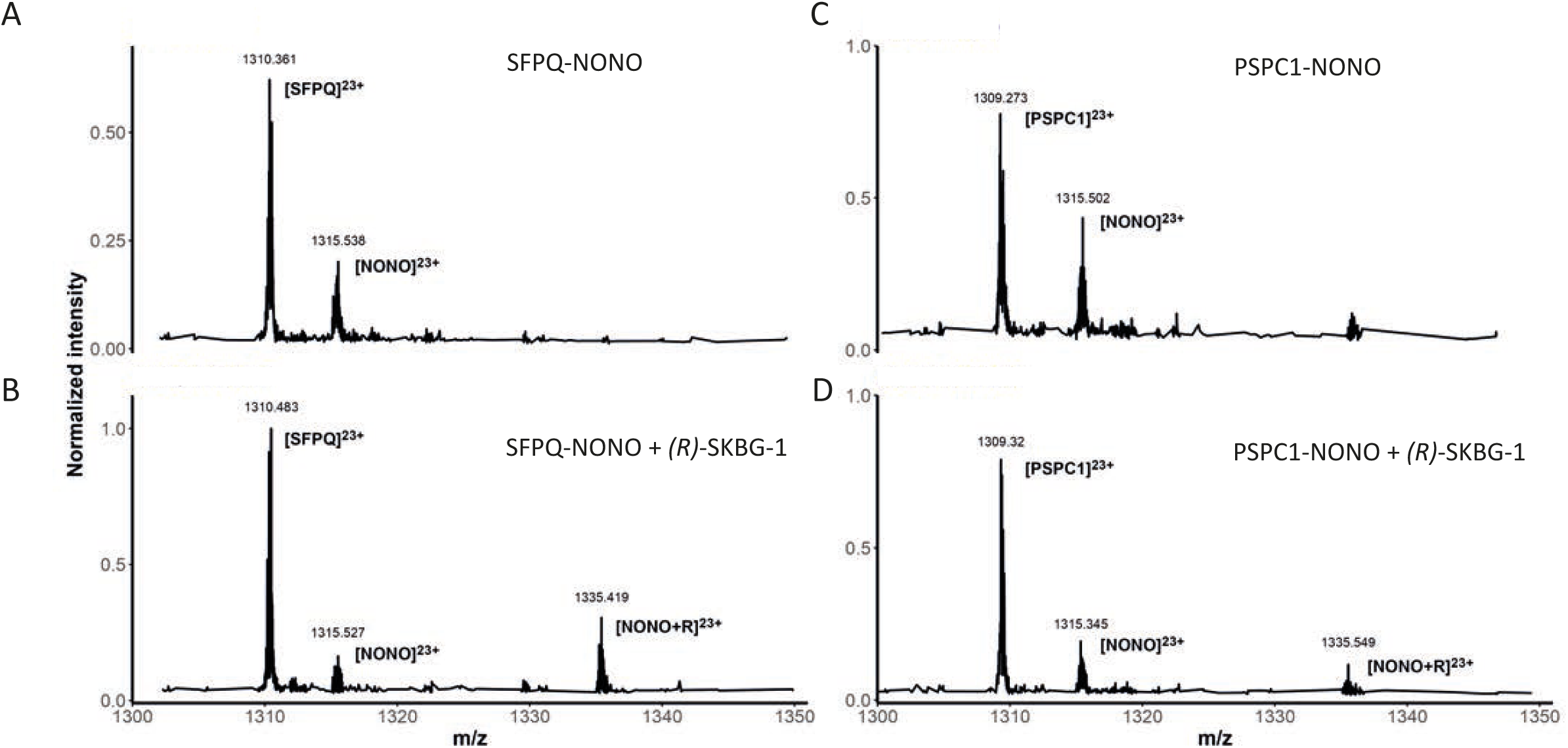
ESI-MS of NONO-PSCP1 and NONO-SFPQ heterodimers before and after treatment with *(R)*-SKBGR-1. A) ESI-MS measurement of the heterodimer of SFPQ(276-535)-NONO(53-312) before (B) and after 2h incubation with *(R)-*SKBG-1. A mass increase that can be attributed to the covalent adduct of (*R*)-SKBG-1 is observed only for NONO and not for SFPQ. Here the most intense charge state (z= +23) is shown. C) ESI-MS measurement of the heterodimer of PSPC1(62-320)-NONO(53-312) before (C) and after 2h incubation with *(R)*-SKBG-1 (D). A mass increase that can be attributed to the covalent adduct of (*R*)-SKBG-1 is observed only for NONO and not for PSPC1. The peak shift measured corresponds to NONO modification by *(R)*-SKBG-1. Full spectra are shown in Fig. S4.

### Structure of NONO homodimer in complex with *(R)-*SKBG-1

In order to determine the molecular determinants driving the selective interaction of NONO with *(R)*-SKBG-1, we set-up crystallization trials of NONO protein covalently modified by *(R)-* SKBG-1 in the presence and in the absence of an RNA oligonucleotide. To do so, the purified NONO(53-312) protein was first incubated with a 2:1 molecular ratio excess of ligand:protein for 2h at 25 °C. The modified protein sample was then concentrated and diluted 3 times with 25 mM Tris-HCl pH 7.5, 250 mM KCl, 0.5 mM EDTA, 50 mM L-Pro to reach a concentration of 200 μM. Commercial crystallization screens were used and protein crystals were harvested and cryo-protected (see Material & methods). X-ray diffraction data were collected on PX1 and PX2A beamlines at the synchrotron SOLEIL. The protein crystallized in *P*4_3_2_1_2 and *P*2_1_2_1_2_1_ space groups containing respectively 1 and 3 homodimers per asymmetric unit. The structures were solved by molecular replacement using NONO atomic coordinates as a model (PDB code 5IFM). The structures were refined to Rfree and Rfactor of 20.3 %/ 27.1 % and 20.6 %/ 26.0 % at 2.9 Å and 2.5 Å resolution respectively (Table S2).

In the crystal n°1 (*P*4_3_2_1_2 space group), the 2 protomers are equivalent and each protomer has a *(R)-*SKBG-1 covalently bound molecule (Fig. 3A). The electron density around the ligand is better defined for chain A than B. Ligand-modified NONO harbors the same fold as the apo NONO with a r.m.s.d of 1.51 Å. In crystal n°2 (in *P*2_1_2_1_2_1_ in the absence of RNA), the asymmetric unit contains three homodimers. The ligand is best defined in chain A, D and partially in F but could not be unambiguously modelled. No ligand could be placed in the other protomers.

**Figure 3:**
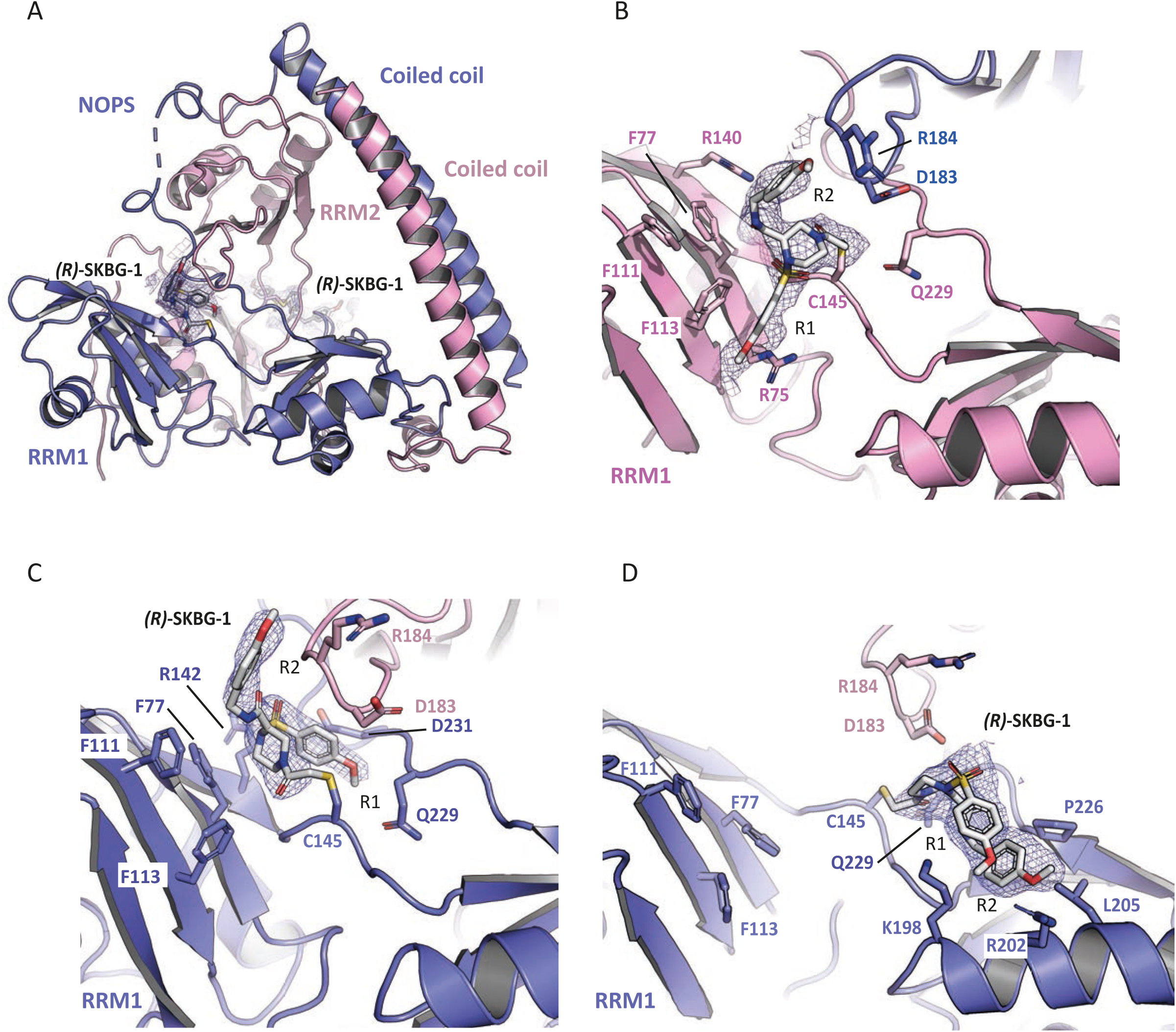
Structure of NONO bound to *(R)*-SKBG-1. A) Overall structure of NONO homodimer covalently modified complex with *(R)-*SKBG-1 (crystal n°1). Each protomer is shown in pink and blue. The composite electron density map contoured at 2.0 σ is shown as a blue mesh around each ligand. Each visible domain is labeled accordingly to the protomer it belongs to. Only one homodimer over the two in the asymmetric unit is shown. B) Close-up view around residue C145 in crystal n°1 (chain B). The composite omit map electron density contoured at 2.0 σ shows the bound ligand on C145 of NONO. Each protomer in colored in blue and pink. C) Close-up view around residue C145 of chain A in crystal n°1 (blue). The electron density of a composite omit map contoured at 2.0 σ shows the bound ligand on residue C145 of NONO. The two protomers are shown in pink and blue. D) Close-up view around residue C145 (chain A) of crystal n°2. (R)-SKBG-1 adopts a “closed” conformation where the p-methoxy-benzene groups are arranged in a close ν-ν interaction. Panel B to D are shown under close orientation in order to illustrate the diversity of ligand orientations observed.

In crystal n°1, the *(R*)-SKBG-1 is bound at C145 residue as expected from previous studies and shown by our *in vitro* experiments ^2^(Fig. 3B). The orientation of the ligand is maintained via hydrophobic interactions provided by the environment at the β-sheet of RRM1 from the first protomer (pink) and residues F77, F111 and F113. Additional polar interactions provided by R140 and Q229 of protomer 1, maintain both R1 and R2 p-methoxy-benzene into a defined conformation. In the other ligand orientation (chain A), *(R*)-SKBG-1 is bound at C145 with the α-chloroacetamide moiety stacking with Phe77 (Fig. 3C). The p-methoxy-benzene group R2 interacts with R184 of the other protomer (pink) while the R1 moiety contacts D142, D231 and Q229 (Fig. 3C).

In crystal n°2 (chain A), the ligand adopts a “closed” conformation where the two p-methoxy-benzene groups stack on each other in a configuration close to ν-ν stacking but with the aromatic rings not harboring a perfect parallel orientation (Fig. 3D). Additional contacts around p-methoxy-benzene group R1 involves K198. The p-methoxy-benzene group R2 contacts R202, L205 and P226. Finally, the α-chloroacetamide moiety is next to Gln229 (same protomer, in blue) and Asp183 of the other protomer (pink).

The heterogeneity in ligand electron density observed for crystal n°2 is highly dependent on the crystal lattice environment. Indeed, there is a correlation with the absence of clear density for the ligand with a poorly crowded environment. In contrast, the good definition observed for crystal n°1 is also correlated with highly ordered environment contributed by symmetry related molecules (Fig. S5A). The other orientation for *(R)-*SKBG-1 in crystal n°2 is depicted in Figure S5B.

### *(R)-*SKBG-1 adopts different conformations on NONO

We have solved the structure of NONO bound to *(R)-*SKBG-1 in two different space groups containing respectively 1 and 3 non-equivalent homodimers. This provides the opportunity to have a combinatorial overview of ligand conformation at its binding site (Fig. 4). We superimposed every homodimer with a bound-ligand to compare the different orientations that *(R)-*SKBG-1 adopts at its binding site. The overall main chain has a good superimposition with a r.m.s.d. <1.5 Å. In three of the observed orientations, the conformations of the bound *(R)-*SKBG-1 are similar with the two p-methoxybenzene groups sandwiched between the β-sheet of the RRM1 of protomer 1 (blue) and the mobile loop corresponding to residues 180 to 185 from the RRM2 of protomer 2 (pink in Fig. 4C & 4D). In the fourth orientation observed in crystal n°2, *(R)-*SKBG-1 adopts a conformation strongly different from the other ones as shown in Fig. 4B. The p-methoxy-benzene rings of *(R)-*SKBG-1 adopt either an extended orientation or a compact orientation via ν-ν stacking.

**Figure 4:**
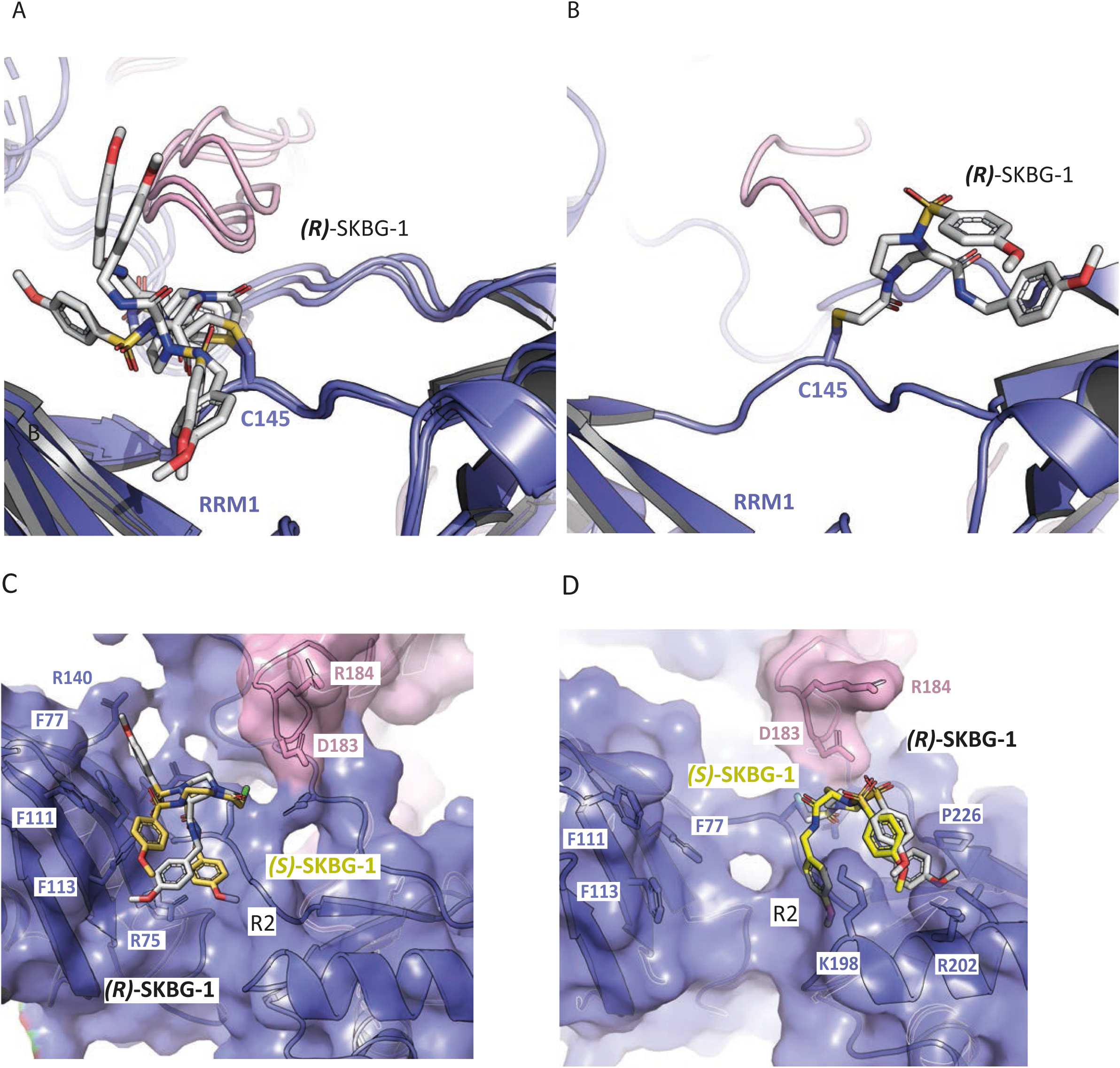
*(R*)-SKBG-1 conformations and enantiomer selectivity. The different conformations of *(R)-*SKBG-1 bound to NONO in our crystal structures are shown. The protein backbone aligns with a r.m.s.d. <1.5 Å and the ligand is displayed. In the different conformations observed, the ligand lies in between the β-sheet of RRM1 of one protomer (blue) and the loop comprised between residues 180-185 of RRM2 of the other protomer (pink) providing additional interactions to stabilize the ligand (A). In the other orientation (B), the ligand is switched by 180° around the C145-α-chloroacetamide bond and lies in an opposite direction. C) *(S)-*SKBG-1 (yellow) was manually superimposed to NONO-bound *(R)-*SKBG-1 (grey) for the main orientations observed in the crystal structures. We used the need of a covalent bond between C145 and the carbonyl moiety of the α-chloroacetamide as a constraint. For each orientation, a sterical clash is observed between NONO and one of the p-methoxy-benzene ring of SKBG-1 compounds. In A) chain D of crystal n°1 (as of Fig. 4A), in D) chain A of crystal n°2 (as of Fig. 4B).

Altogether this suggests that, apart from the covalent link with C145, *(R*)-SKBG-1 has a large degree of freedom to adapt to its local environment. This is also in agreement with the poor definition at other C145 sites in other chains in crystal n°1 and crystal n°2 (see Discussion).

## DISCUSSION

### A mobile covalently-bound ligand

Covalently modified NONO has been crystallized in two different space groups with one (crystal n°1) and three (crystal n°2) non-equivalent homodimers in the asymmetric unit respectively. Structural alignment of the protein backbone reveals that *(R)*-SKBG-1 can adopt different conformation when covalently bound to the protein (Fig. 4).

In the two crystal forms, the environment around the C145 residue targeted by the *(R)-*SKBG-1 is different. Since the covalent attachment of the *(R)-*SKBG-1 is very efficient according to our ESI-MS data, and since we used the same batch of modified protein sample to perform crystallization and ESI-MS, the absence of a corresponding electronic density near some targeted cysteines is likely due to a high mobility of the ligand and C145 proximal environment. Indeed, the differences in crystal packing promotes the observation of a portfolio of conformations for *(R)-*SKBG-1. When the local environment around C145 is open, the *(R)-*SKBG-1 is highly flexible and we do not observe significant electronic density preventing ligand modelling. However, when the protein chains are more tightly packed due to the crystal lattice as shown Fig. S5A, the electron density around the *(R)-*SKBG-1 is well defined. From that study, it appears the ligand biological effect is likely due to the combination of two characteristics of the ligand, (i) its ability to a make covalent bond with the C145. This provides specificity over the two other DBHS proteins SFPQ and PSPC1 since there is no cysteine residue at the equivalent position in SFPQ and PSPC1; (ii) the ability of the ligand to adapt to its local environment due to a high degree of flexibility.

### Structural basis for NONO’s enantiomer selectivity

NONO displays enantiomer selectivity for *(R)-*SKBG-1 over *(S)-*SKBG-1. This was reported from the initial publication ^2^ and confirmed by ESI-MS analysis in our *in vitro* study. Very limited modification of NONO is observed for *(S)*-SKBG-1 under the identical conditions in comparison to *(R)-*SKBG-1 (Fig. 1). The crystal structures of *(R)-*SKBG-1-bound NONO in the two main orientations observed and its comparison with *(S)-*SKBG-1 gives a rationale for such selectivity (Fig. 4C, 4D and Fig. S6). Despite the ability of the p-methoxy-benzene rings to rotate around their covalent bond with the α-chloroacetamide, the (*S*)-configuration of the α-chloroacetamide (yellow in Fig. 4) forces a ring orientation that apparently clashes with the protein. This is the case for the orientation when *(R*)-SKBG-1 packs at the RRM1 β-sheet (Fig. 4C) where R2 p-methoxybenzene ring of *(S)*-SKBG-1 would clash with residues next to C145. The presence of the oxygen atom from the amide moiety forces the rotation of the R2 ring, orienting the 2 p-methoxy-benzene directly towards the protein, a situation which is not found in the (*R*) enantiomer. A similar conclusion can be drawn from the superimposition of *(S)-*SKBG-1 in the opposite orientation (Fig. 4D) where the R2 p-methoxy benzene ring orientation is forced to clash with at least K198 in the (*S*) enantiomer.

In both structures, the experimental orientation is largely constrained by crystal contacts and is thus not reflecting a physiological situation where molecular crowding is very different. Our experimental observations (ESI-MS and experimental models) argue for no possibility to engage the *(S)*-SKBGR-1 into a covalent bond formation.

Future developments of NONO-specific compounds may benefit from these first experimental models. In particular, these data may help explain how chemical modifications of the benzene substituent groups lead to compounds with varying affinities, as observed by Kathman (Kathman et al., 2023). The comparison of B21 with compounds 8 and 9 provides valuable insights. Specifically, the presence of a meta-positioned electron-donating methyl group on the sulfonamide-linked benzene ring (R1) in compound 8, or a protected amide function on R2 in compound 9, results in a loss of affinity. Further structural modifications of the R1 and R2 motifs, especially through the introduction of polar groups capable of hydrogen bonding, could improve ligand anchoring to the protein.

### Impact of *(R)-*SKBGR-1 covalent binding on RNA interaction

It has been reported that covalent modification of NONO by *(R)-*SKBG-1 enhances its RNA binding properties leading to its accumulation in RNA foci ^2^. Even though crystal n°1 has been obtained in the presence of an RNA oligonucleotide in the crystallization condition, no density can be attributed to the RNA. Crystal packing around *(R)-*SKBG-1 is such that no free space is left to accommodate an RNA molecule. Hence, despite our efforts we have not been able to obtain a crystal structure of NONO in a ternary complex with an RNA and *(R)-*SKBG*-*1. In the absence of a structure of NONO in complex with an RNA and given our experimental observations that *(R)-*SKBG-1 has a large conformational repertoire upon NONO binding, it is difficult to propose a conclusive model for RNA binding by NONO in the presence of a ligand at this stage. The use of the recently and only reported structure of a DBHS protein (SFPQ) in complex with an RNA (PDB 7UJ1)^14^ can be used to analyze how *(R)-*SKBG-1 may enhance RNA binding. This model was superimposed onto the *(R)-*SKBG-1 NONO homodimer for each of the four observed orientations of *(R)-*SKBG-1. The two other orientations of *(R)-*SKBG-1 are *per se* compatible with RNA binding with no observed clash (Fig. S7A & 7B). Two orientations would be compatible with RNA binding, providing local rearrangements of side chains, RNA bases and ligand (Fig. S7C and 7D).

At this stage such modelling has the advantage of giving an initial template for the design of novel ligands. It is likely that the p-methoxy-benzene rings play a role in stacking RNA bases to the protein. In the absence of an RNA-bound ligand-modified NONO, further experimental structures with different ligands are necessary to optimize or to rationalize the observed *in vivo* effects on RNA binding enhancement.

## ACKNOWLEDGEMENTS

This work was funded by the “Ligue Régionale Contre le Cancer” Comité de Charente-Maritime (CD17) et Comité des Landes (CD40), INSERM, CNRS and the University of Bordeaux. This work benefited from access to Plateforme de BioPhysico-Chimie Structurale of the IECB (Univ. Bordeaux, CNRS UAR3033, Inserm US001) for mass spectrometry.

We acknowledge SOLEIL for provision of synchrotron radiation facilities and we would like to thank Pierre Montaville, Pierre Legrand, William Shepard for assistance in using beamlines PX1 and PX2A.

We also thank B.F Cravatt, Dr Melillo for kindly providing us with *(R)-*SKBG-1 and *(S)-*SKBG-1. We thank Drs S. Thore, C.D. Mackereth and V. Desvergnes for critical reading of the manuscript and fruitful discussions. We thank Drs Matthieu Ranz and Drs V. Desvergnes for their help in generating ESI-MS figures and Fig. S2.

## DATA AVAILABILITY

The crystal structures were deposited in the RCSB Protein Data Bank under accession codes 9QZL and 9QXZ.

## AUTHORS’ CONTRIBUTION

AVF was supported by the ERASMUS mundi program.

AVF and SF purified the proteins and prepared the samples.

CB performed the ESI-MS analysis.

SF supervised the work and wrote the article.

## STAR METHODS

### Molecular biology

The ORF encoding hNONO(53-312), hPSPC1(62-320) and hSFPQ(276-535) were cloned into a pET-15b derived plasmid allowing for protein expression in frame with an N-terminal hexahistidine tag and a TEV cleavage site, and NONO(53-312) was also cloned into a pCDF derived plasmid allowing protein expression of an untagged protein. Point mutagenesis of hNONO C145V was obtained by PCR and Gibson assembly cloning and further confirmed by DNA sequencing.

### Protein expression and purification

Protein expression of NONO(53-312), SFPQ(276-535)-NONO(53-312) and PSPC1(62-320)-NONO(53-312) was achieved by overexpression in Rosetta2 cells. Electrocompetent cells were transformed with the required plasmid(s) and plated onto LB-agar plates supplemented with Ampicilin at 100 μg/ml and Chloramphenicol 34 μg/ml and, when necessary (pCDF plasmid), with 50 μg/ml Streptomycin. 1L culture in TB was then inoculated and grown at 37 °C up to an OD 600 of > 2.0. After 2 h at 15 °C, the protein expression was induced overnight with 0.25 mM IPTG. Cells were harvested the next day by centrifugation at 4500 g for 15 min. Cell pellet was resuspended in TB media and sonicated for 3 min at 7W under spinning. After 45 min centrifugation at 50,000 g at 4 °C, the supernatant was incubated with cobalt-affinity resin at 4 °C for 30 min. Resin was then packed in a AK10/26 column and extensively washed with 25 mM Tris-HCl pH 7.4, 1000 mM KCl then 25 mM Tris-HCl pH 7.4, 150 mM KCl. Elution was achieved by a gradient of imidazole up to 250 mM in 25 mM Tris-HCl pH 7.4, 150 mM KCl. The peak was collected and supplemented with 1 mM DTT, 0.5 mM EDTA and incubated at 16 °C overnight with TEV protease at a ratio 1:100 w/w. The protein sample was then applied onto a Heparin column equilibrated in 25 mM Tris-HCl pH 7.4, 100 mM KCl. After washes in 25 mM Tris-HCl pH 7.4, 100 mM KCl, the protein sample was eluted with a gradient of KCl up to 1000 mM. Bound protein was harvested, concentrated and exchanged for a buffer containing 25 mM Tris-HCl pH 7.4, 500 mM KCl, 10 % w/v.

The theoretical mass of each purified protein is summarized in Table 1. The difference in size for NONO is associated to the presence of Gly-His residues at the N-terminus after TEV cleavage.

### Protein modification

Proteins were kept in 25 mM Tris-HCl pH 7.4, 500 mM KCl, 10 % w/v at 100 μM. In order to ensure a complete chemical modification assuming one specific site per monomer, proteins were mixed with an mild excess of ligand at 2:1 molecular ratio (ligand:protein monomer) and incubated at 25 °C for 2h. The excess of ligand was removed by iterative dilution/concentration steps with an amicon ultrafiltration 30kDa with a buffer containing 25 mM Tris-HCl pH 7.5, 250 mM KCl, 0.5 mM EDTA, 50 mM L-Pro. The same samples were lately used for ESI-MS and protein crystallization.

### Complex preparation and analysis by ESI-MS

In order to analyze the various sample by ESI-MS, protein sample buffer was exchanged for ammonium acetate 500 mM pH 7.4 by iterative dilution/concentration steps (at least 5 times) by adding 15 ml of buffer to < 1ml of sample between each round of centrifugation. Then, samples were diluted in 50/50 CH_3_OH/H_2_O + 1% formic acid in order to obtain approximately a concentration of 2 µM.

Mass spectra were acquired with a Bruker SolariX 7 T XR Fourier transform ion cyclotron resonance mass spectrometer (Bruker Daltonics, Bremen, Germany). The capillary voltage was set at 4.5 kV in the positive ion mode. The drying-gas temperature was 200 °C and its flow rate 4 L/min. The ion-funnel RF amplitude was 150 Vpp, the funnel 1 voltage was 150 V and the skimmer 1 voltage was 100V. RF frequencies used were the following: octopole (2 MHz), quadrupole (2 MHz) and transfer line (4 MHz). The time-of-flight was 1.2 ms and Q1 Mass was set at 400 m/z. The source region (PS1) pressure was 3.9 mbar; the quadrupole region (PS4) pressure was 7.0 × 10^-6^ mbar, and the trap-chamber pressure (PS6) was 2.4 × 10^-10^ mbar. Fifty scans were averaged for each spectrum. External calibration was done by infusing the ESI-L calibrant (Low concentration tuning mix, ref G1969-85000, Agilent). The protein sample was delivered by the syringe pump incorporated to the FTICR mass spectrometer at flow rate 3 µL/min. Spectra were extracted using Data Analysis (Bruker Daltonics) software and revised manually.

### Protein crystallization and structure determination

NONO(53-312) was modified with a protein:ligand of 1:2 molecular ratio for 2h at 25°C in 20 mM Tris-HCl pH 7.5, 250 mM KCl, 0.5 mM EDTA, 50 mM L-Pro buffer. The excess of ligand was then removed by a series of 3 concentration and dilution steps with buffer in 20 mM Tris-HCl pH 7.5, 250 mM KCl, 0.5 mM EDTA, 50 mM L-Pro buffer to reach a final concentration of 3 mg/ml (around 200 μM). The modified protein was then crystallized using commercial screens in the presence and in the absence of an RNA oligonucleotide (AAAAAGGUAAG). Best crystals were found in Index G8 (in the absence of RNA) and G9 conditions (in the presence of RNA) (0.2 M Ammonium acetate, 0.1 M HEPES pH 7.5 or Tris-HCl pH 8.5, 25 % w/v PEG 3350). Crystals were transferred in a cryo buffer containing 35 % w/v PEG 3350 and flash-frozen in liquid nitrogen. Diffraction data were collected at SOLEIL PX1 and PX2A beamlines. Data were then reduced and SCALED using the last version of XDS ^15,16^. The structure was solved by molecular replacement in Phenix suite using one dimer of NONO(53-312)(PDB= 5IFM)^17^. The asymmetric unit comprised 3 dimers of NONO that were refined by successive iterations with Buster 2.11.2 ^18^ and manual fitting in Coot ^19^. The ligand restraints were generated with Grade2 (https://grade.globalphasing.org/cgi-bin/grade2_server.cgi). The last steps of refinement were devoid of crystallographic symmetry to avoid bias in ligand positioning. The final models were refined to a Rfree/ Rfac of 19.1 %/ 27.7 % and 20.8 %/ 26.0 % respectively (Table S1).

## Supplementary Data

**Table S1:** Average theoretical masses of protein constructs.

|  | Mass (in Da) |
| --- | --- |
| **NONO(53-312)(post TEV cleavage)** | **30,425.20** |
| **NONO(53-312)(native)** | **30,231.62** |
| **SFPQ(276-535) (post TEV cleavage)** | **30,115.10** |
| **PSPC1(62-320) (post TEV cleavage)** | **30,092.29** |

**Table S2:**
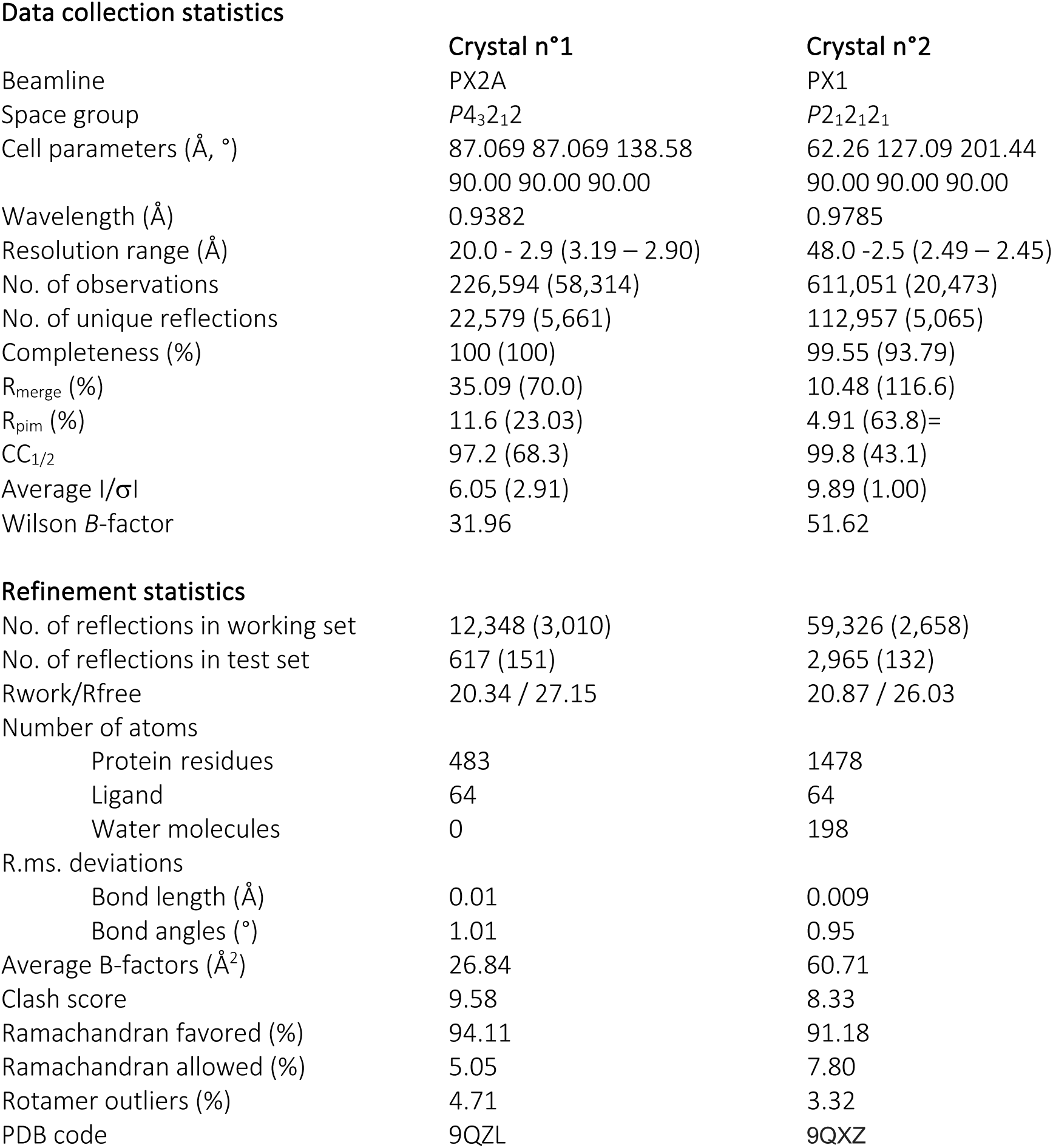
Crystal data, diffraction data collection and refinement statistics.

Data collection statistics
|  | Crystal n°1 | Crystal n°2 |
| --- | --- | --- |
| Beamline | PX2A | PX1 |
| Space group | $P4_32_12$ | $P2_12_12_1$ |
| Cell parameters (Å, °) | 87.069 87.069 138.58<br>90.00 90.00 90.00 | 62.26 127.09 201.44<br>90.00 90.00 90.00 |
| Wavelength (Å) | 0.9382 | 0.9785 |
| Resolution range (Å) | 20.0 - 2.9 (3.19 – 2.90) | 48.0 -2.5 (2.49 – 2.45) |
| No. of observations | 226,594 (58,314) | 611,051 (20,473) |
| No. of unique reflections | 22,579 (5,661) | 112,957 (5,065) |
| Completeness (%) | 100 (100) | 99.55 (93.79) |
| $R_{\text{merge}}$ (%) | 35.09 (70.0) | 10.48 (116.6) |
| $R_{\text{pim}}$ (%) | 11.6 (23.03) | 4.91 (63.8)= |
| $CC_{1/2}$ | 97.2 (68.3) | 99.8 (43.1) |
| Average $I/\sigma I$ | 6.05 (2.91) | 9.89 (1.00) |
| Wilson $B$-factor | 31.96 | 51.62 |
| Refinement statistics |  |  |
| No. of reflections in working set | 12,348 (3,010) | 59,326 (2,658) |
| No. of reflections in test set | 617 (151) | 2,965 (132) |
| Rwork/Rfree | 20.34 / 27.15 | 20.87 / 26.03 |
| Number of atoms |  |  |
| Protein residues | 483 | 1478 |
| Ligand | 64 | 64 |
| Water molecules | 0 | 198 |
| R.ms. deviations |  |  |
| Bond length (Å) | 0.01 | 0.009 |
| Bond angles (°) | 1.01 | 0.95 |
| Average B-factors (Å<sup>2</sup>) | 26.84 | 60.71 |
| Clash score | 9.58 | 8.33 |
| Ramachandran favored (%) | 94.11 | 91.18 |
| Ramachandran allowed (%) | 5.05 | 7.80 |
| Rotamer outliers (%) | 4.71 | 3.32 |
| PDB code | 9QZL | 9QXZ |

**Figure S1:**
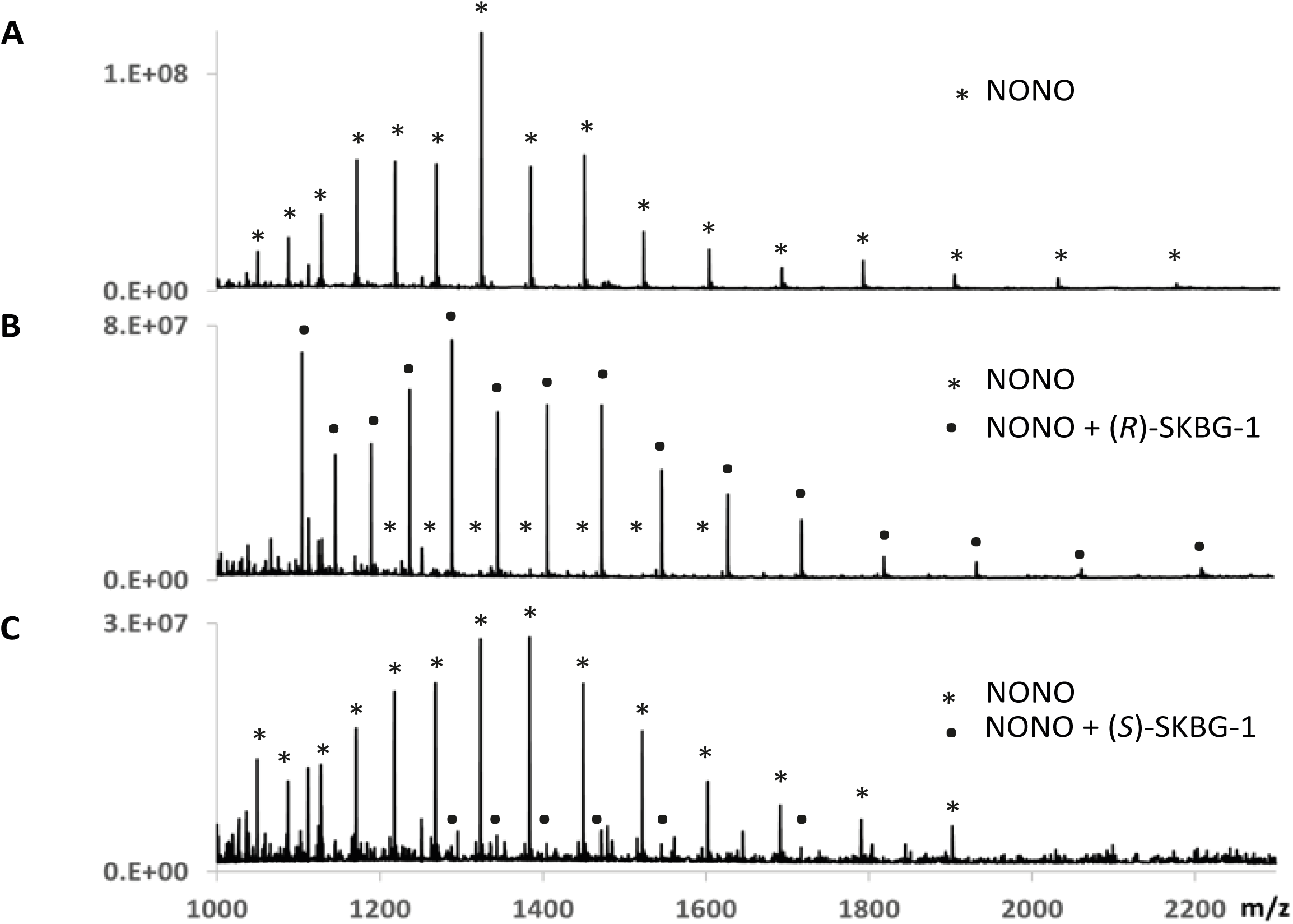
Full ESI-MS spectra corresponding to NONO(53-312) before (A) and after modification with (*R*)-SKBG-1 (B) and (*S*)-SKBG-1 (C). Peak corresponding to native NONO are marked with (*) and to ligand-bound NONO with (•).

**Figure S2:**
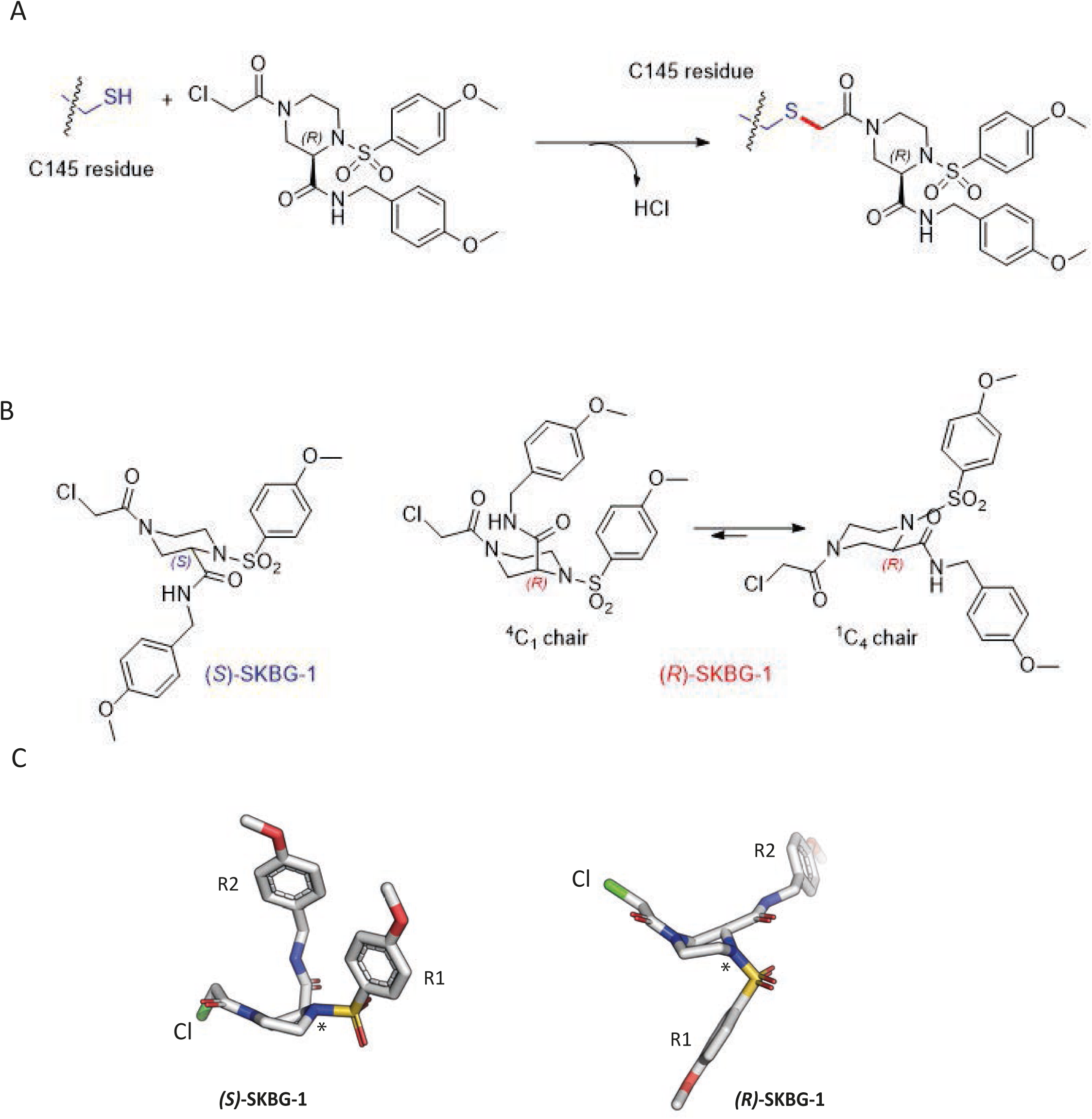
2D and 3D representations of (*R*)-SKBG-1 and (*S*)-SKBG-1 and the chemical reaction with NONO C145 leading to the covalent adduct. A) The thiol group of cysteine C145 reacts with the ligand via an SN2 nucleophilic substitution, formally resulting in the loss of HCl and the formation of a new covalent S–C bond. B) 2D representation of *(S*)-SKBG-1 (left) and *(R*)-SKBG-1 (right). C) 3D representation of *(S*)-SKBG-1 (left) and of *(R*)-SKBG-1 (right). The chlorine leaving atom is shown in green. The chiral center is marked with a “*”.

**Figure S3:**
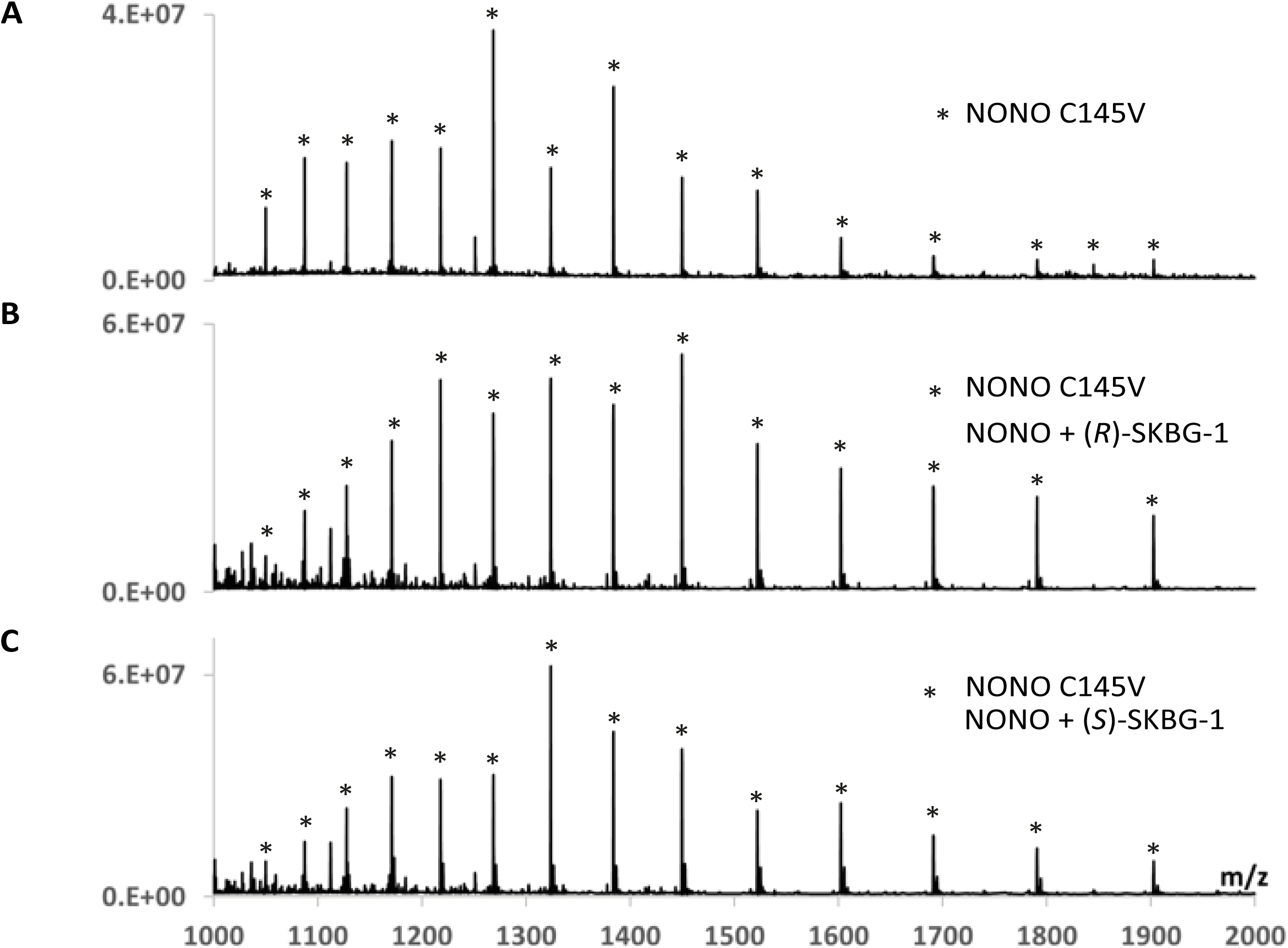
Full ESI-MS spectra corresponding to NONO(53-312) C145V before (A) and after modification with (*R*)-SKBG-1 (B) and (*S*)-SKBG-1 (C). Peak corresponding to native NONO C145V are marked with (*).

**Figure S4:**
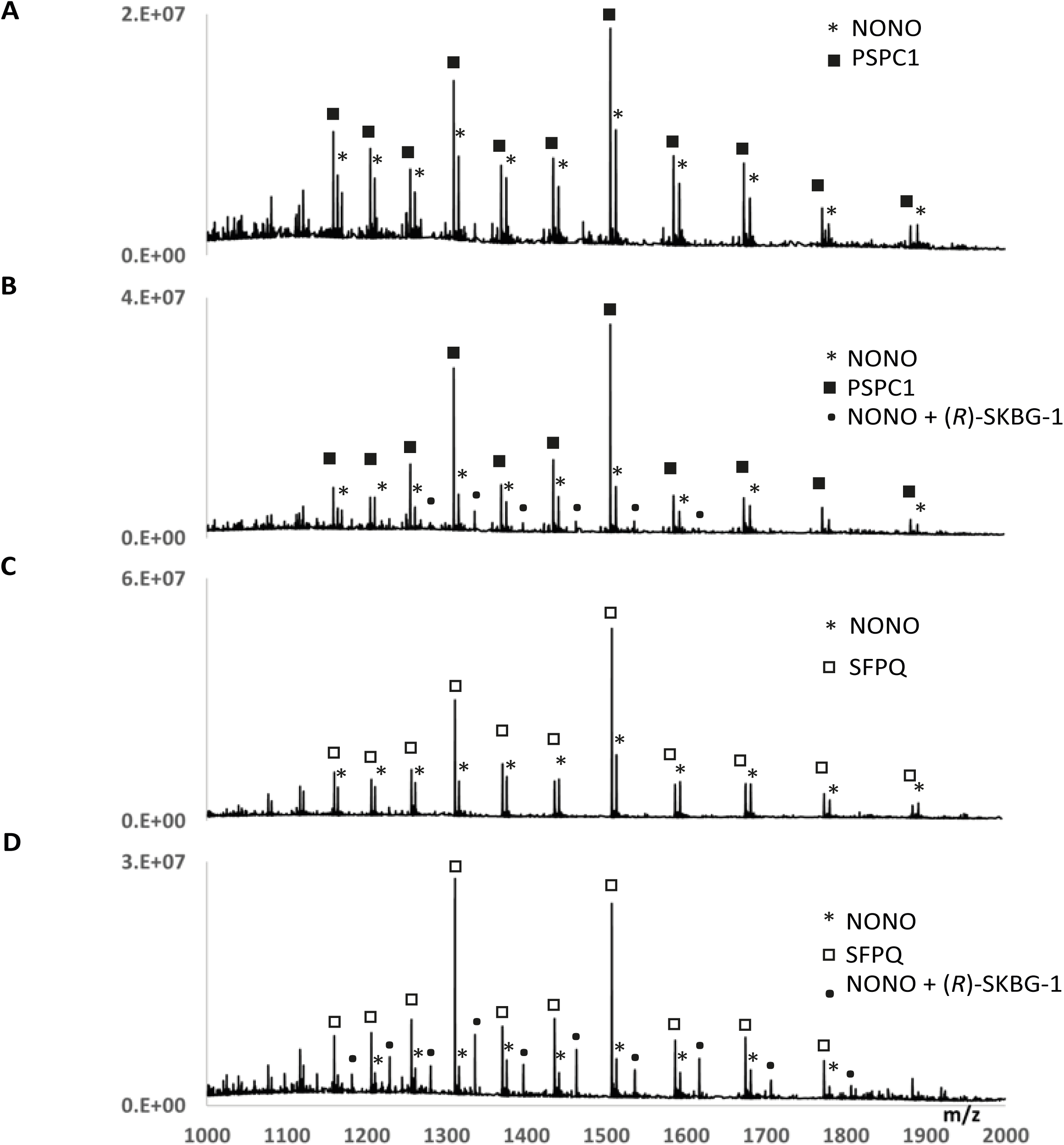
Full ESI-MS spectra corresponding to NONO(53-312)-PSPC1(62-320) and NONO(53-312)-SFPQ(276-535) before (A and C) and after modification with (*R*)-SKBG-1 (B and D) respectively. Peak corresponding to native NONO are marked with (*) and to ligand-bound NONO with (•). Peak corresponding to PSPC1 and SFPQ are labelled (▪) and (□) respectively.

**Figure S5:**
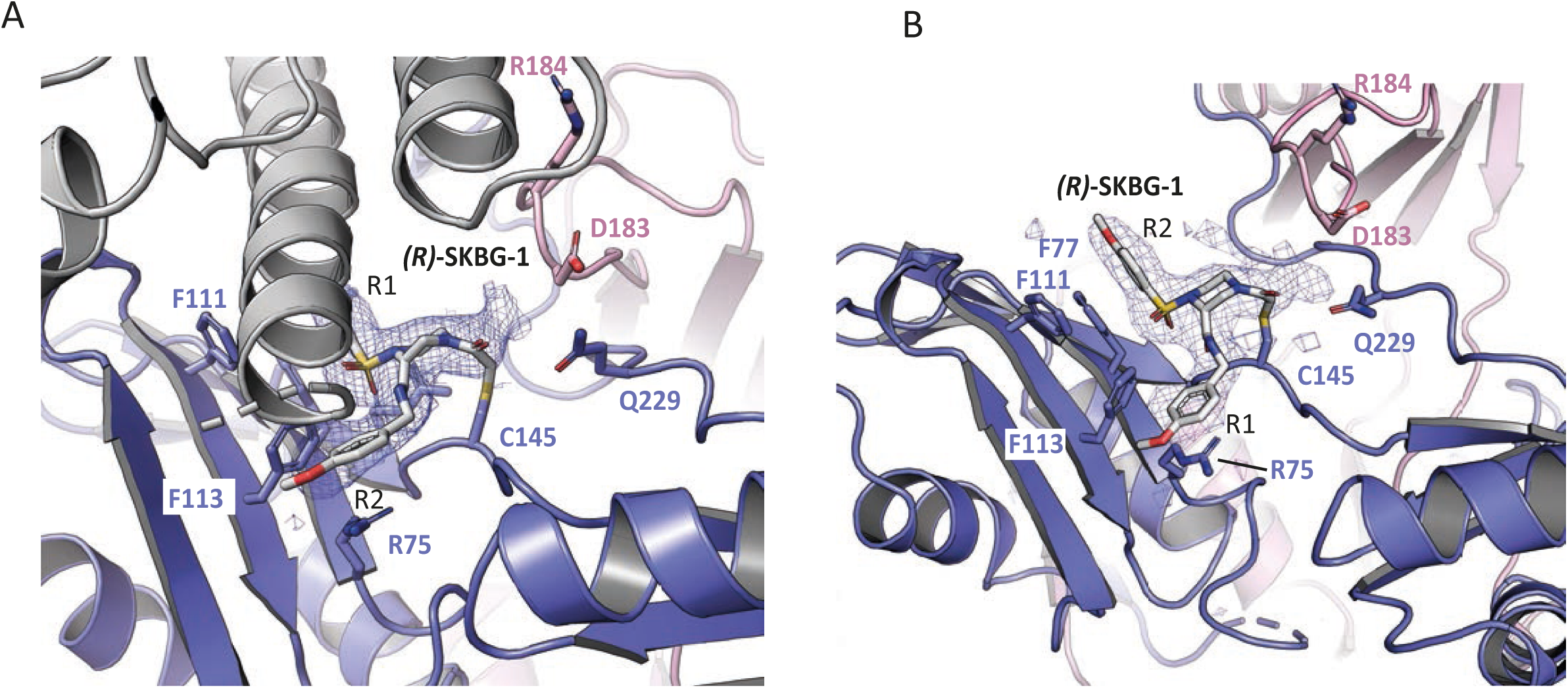
A) Composite omit map contoured at 4σ around the ligand binding site as shown in Figure 4B is displayed. The coiled coil domain of a symmetry related molecule (grey) packs along the ligand. B) Composite omit map contoured at 4σ around the ligand binding site of chain D in crystal n°2. *(R*)-SKBG-1 packs via R2 ring with F77, while F113 packs with R1 ring. Additional contacts are provided by R75.

**Figure S6:**
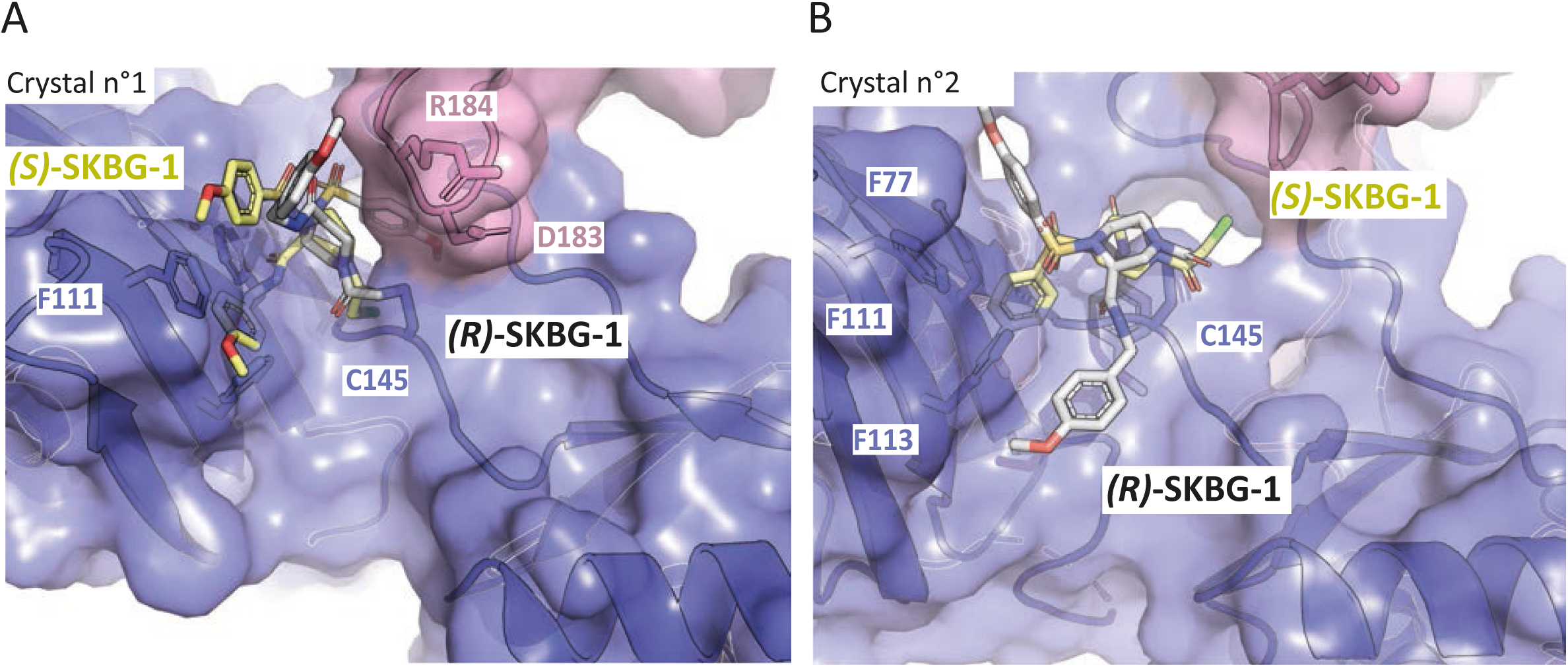
Enantiomer selectivity of NONO for *(R)-*SKBG-1 versus *(S)-*SKBG-1. *(S)-*SKBG-1 (yellow) was manually superimposed to NONO-bound *(R)-*SKBG-1 (grey) for the main orientations observed in the crystal structures. We used the need of a covalent bond between C145 and the carbonyl moiety of the α-chloroacetamide as a constraint. For each orientation, a sterical clash is observed between NONO and one of the p-methoxy-benzene ring of *(S)*-SKBG-1 compounds. Here are displayed the three other orientations not shown in the main text. In A) chain A of crystal n°1, in B) chain D of crystal n°2.

**Figure S7:**
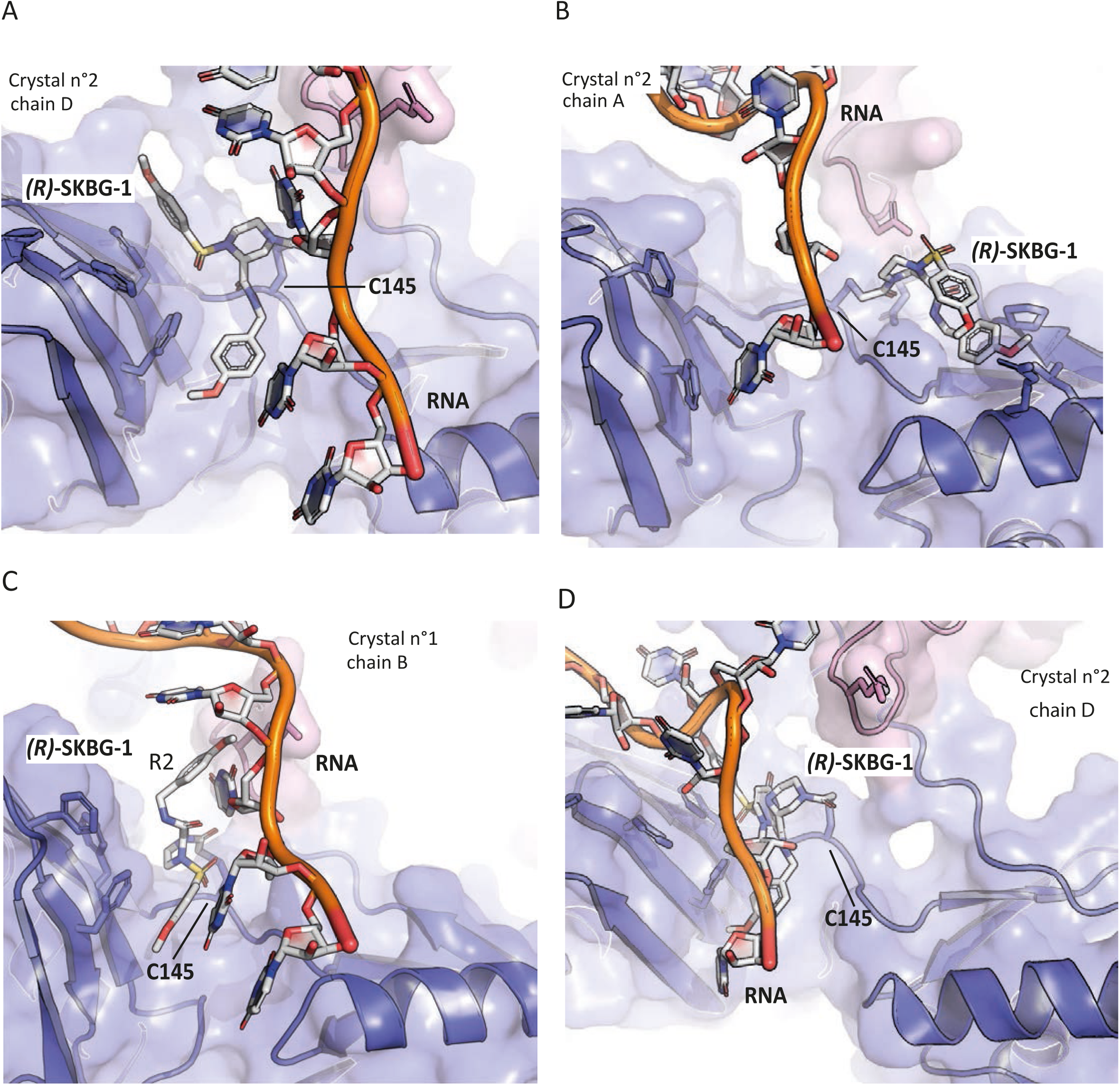
Model of a ternary complex of NONO bound to *(R)-*SKBG-1 and RNA. The SFPQ-RNA model structure (PDB 7UJ1)(Wang *et al*, 2022) has been used to generate a model of RNA-bound NONO in the presence of *(R)-*SKBG-1. The SFPQ ribbon representation is omitted to keep only the RNA model. Two out of the four observed orientations present no clash between the ligand and the RNA (S7A and S7B). Two orientations would clash with the RNA binding if no reorientation of the ligand and/or the RNA bases would not happen (S7C and S7D). A) Crystal n°2, Chain D B) Crystal n°2, Chain A C) Crystal n°1, Chain B D) Crystal n°2, Chain D

